# Artificial intelligence based computational framework for drug-target prioritization and inference of novel repositionable drugs for Alzheimer’s disease

**DOI:** 10.1101/2020.07.17.208116

**Authors:** Shingo Tsuji, Takeshi Hase, Ayako Yachie, Taiko Nishino, Samik Ghosh, Masataka Kikuchi, Kazuro Shimokawa, Hiroyuki Aburatani, Hiroaki Kitano, Hiroshi Tanaka

**Affiliations:** Research Center for Advanced Science and Technology, The University of Tokyo, 4-6-1 Komaba, Meguro-ku, Tokyo 153-8904 JAPAN; The Systems Biology Institute, Saisei Ikedayama Bldg. 5-10-25 Higashi Gotanda Shinagawa, Tokyo 141-0022, Japan; Medical Data Sciences office, Tokyo Medical and Dental University, 20F, M&D Tower, 1-5-45 Yushima, Bunkyo-ku, Tokyo 113 - 8510, JAPAN; Department of Genome Informatics, Graduate School of Medicine, Osaka University, 2-2 Yamadaoka, Suita, Osaka 565-0871, Japan; Center for Mathematical Modeling and Data Science, Osaka University, 1-3 Machikaneyama-cho, Toyonaka City, Osaka 560-8531, Japan

**Keywords:** network bedding, deep learning, machine learning, systems biology, drug discovery, protein interaction network

## Abstract

**Background:** Identification of novel therapeutic targets is a key for successful drug development. However, the cost to experimentally identify therapeutic targets is huge and only 400 genes are targets for FDA-approved drugs. Therefore, it is inevitable to develop powerful computational tools to identify potential novel therapeutic targets. Because proteins make their functions together with their interacting partners, a protein-protein interaction network (PIN) in human could be a useful resource to build computational tools to investigate potential targets for therapeutic drugs. Network embedding methods, especially deep-learning based methods would be useful tools to extract an informative low-dimensional latent space that contains enough information required to fully represent original high-dimensional non-linear data of PINs.

**Results:** In this study, we developed a deep learning based computational framework that extracts low-dimensional latent space embedded in high-dimensional data of the human PIN and uses the features in the latent space (latent features) to infer potential novel targets for therapeutic drugs. We examined the relationships between the latent features and the representative network metrics and found that the network metrics can explain a large number of the latent features, while several latent features do not correlate with all the network metrics. The results indicate that the features are likely to capture information that the representative network metrics can not capture, while the latent features also can capture information obtained from the network metrics. Our computational framework uses the latent features together with state-of-the-art machine learning techniques to infer potential drug target genes. We applied our computational framework to prioritized novel putative target genes for Alzheimer’s disease and successfully identified key genes for potential novel therapeutic targets (e.g., DLG4, EGFR, RAC1, SYK, PTK2B, SOCS1). Furthermore, based on these putative targets, we inferred repositionable candidate-compounds for the disease (e.g., Tamoxifen, Bosutinib, and Dasatinib)

**Discussions:** Our computational framework could be powerful computational tools to efficiently prioritize new therapeutic targets and drug repositioning. It is pertinent to note here that our computational platform is easily applicable to investigate novel potential targets and repositionable compounds for any diseases, especially for rare diseases.

## Background

Biomedical research, especially drug discovery, is now going through a global paradigm shift with AI (Artificial Intelligence) technologies and their application to Big Data in biomedical domain [1–3]. The complex, non-linear, multi-dimensional nature of big data gives us unique challenges and opportunities in their processing and analysis to obtain actionable insights. Particularly, existing statistical techniques like principle components analysis (PCA) are insufficient for capturing the complex interaction patterns hidden in multiple dimensions across the spectrum of data [4]. Thus, a key challenge for future drug discovery research is to develop AI based powerful computational tools that can capture biomedical insights in multiple dimensions and obtain “value#x201D; in the form of actionable insights (e.g., insights towards selecting and prioritizing candidate targets and repositionable drug for the candidate targets) from volume of big data.

“Big Data#x201D; in biomedical domain are generally with high dimensionality. Their dimensionality should be reduced to avoid undesired properties of high-dimensional space, especially the curse of dimensionality [5]. Dimensionality reduction techniques facilitate classification, data visualization, and high-dimensional data compression [6]. However, classical dimensional reduction techniques (e.g., PCA) are generally linear techniques and thus insufficient to handle non-linear data [4, 6].

With a recent advancement of AI technologies, a large number of dimensionality reduction techniques for non-linear complex data are available [4,6,7]. Among dimensionality reduction techniques, a multi-layer neural network based technique, “deep autoencoder#x201D;, could be the most powerful technique to reduce dimensionality of non-linear data [4, 6]. Deep autoencoders composed of multilayer encoder and decoder networks. Multilayer encoder component transforms high-dimensionality of data into low-dimensional representation, while multilayer “decoder” component recovers original high-dimensional data from the low-dimensional representation. Weights associated with links connecting the layers are optimized by minimizing the discrepancy between input and output of the network, i.e., in ideal condition, the values of nodes in input layer is same as those in output layer. After the optimization steps, the middle-hidden encoder layer gives a law dimensional representation that preserves information contained is original data as much as possible [6]. The values of nodes in the middle-hidden encoder layer would be useful features for classification, regression, and data visualization of high-dimensional data.

In drug discovery research, a key for successful development of therapeutic drug is to identify novel drug-targets [8–10]. However, the cost to experimentally predict drug target is huge and only ‘400 genes are used as targets of FDA-approved drugs [11]. Thus, it is inevitable to develop a powerful computational framework that can identify potential novel drug-targets.

PIN data could be a useful big resource to computationally investigate potential novel drug-targets, because proteins make their functions together with their interacting partners and network of protein interaction captures down-stream relationships between targets and proteins [8–10,12]. With a recent advancement of network science, various network metrics are now available and have been used to investigate structure of molecular interaction networks and their relationships with drug-target genes [8–10,12,13]. For example, degree, the number of links to a protein, is a representative network metric to investigate molecular interaction networks [10], i.e., almost all FDA-approved drug-targets are middle- or low-degree proteins, while there are almost no therapeutic targets among high-degree proteins. It indicates that key features for identification of potential drug target genes could be embedded in the complex architectures in the protein-protein interaction networks [10].

Data of genome wide PINs are typical non-linear high-dimensional big-data in biomedical domain and are composed of thousands of proteins and more than ten-thousands of interactions among them [8,9]. Mathematically, a protein-protein interaction network is represented as adjacency matrix [14]. The adjacency matrices for PINs are with rows and columns labelled by proteins and elements in the matrices are represented as binary value, i.e., 1 or 0 in position (*i,j*) according to whether protein *i* interacts with protein *j* or not. In the adjacency matrix, each row represents interacting pattern for each protein and may be useful features to predict potential drug target proteins. However, the feature vector for a protein is high dimensional (e.g., several thousands dimensions) and also sparse, because protein interaction network composed of thousands of proteins and the number of columns (features) of each proteins is very large [14]. In order to use data of PIN effectively for drug discovery research, we need apply powerful dimensional reduction techniques to high-dimensional data of PIN.

Recently, researchers have developed “network embedding” methods that apply dimensional reduction techniques to extract low-dimensional representations of a large network from high-dimensional adjacency matrix of the network [14,15]. For examples, several researchers have used singular value decomposition and non-negative matrix factorization methods to map high-dimensional adjacency matrices of large-scale networks into low-dimensional representations [16,17]. However, the feature vector for a protein is high dimensional (e.g., several thousands dimensions) and also a sparse, because protein interaction network composed of thousands of proteins and the vast majority of proteins in PIN have few interactions [14].

In order to address this issue, several researchers have used network embedding methods based on deep learning techniques [18,19]. Especially, deep autoencoder based network embedding methods would be useful to transform non-linear large-scale networks into low-dimensional representations. Wang et al. applied deep autoencoder based network embedding method to large scale social networks (e.g., arxiv-GrQc, blogcatalog, Flicker, and Yutube) and successfully map these networks on low-dimensional representations [18].

In this study, in order to infer potential novel target genes, we proposed a computational framework based on a representative network embedding method that uses deep autoencoder to map a genome-wide protein interaction network into low-dimensional representations. The framework builds a classifier based on state-of-the-art machine learning techniques to predict potential novel drug-targets using the resultant low-dimensional representations. We applied the framework to predict potential novel drug targets for Alzheimer’s disease. Based on the list of predicted candidate novel drug targets, we further infer potential repositionable drug candidates for Alzheimer’s disease.

## Results and Discussions

In this study, we proposed a computational framework (as show in Figure 1) to predict potential drug target genes using information of genome-wide protein-protein interaction networks. The framework uses a representative network embedding method based on deep autoencoder to extract low-dimensional features for each gene from the PIN. Then, by using the extracted low-dimensional features as training data, the framework builds a machine-learning model to predict potential drug-target proteins.

**Figure 1.**
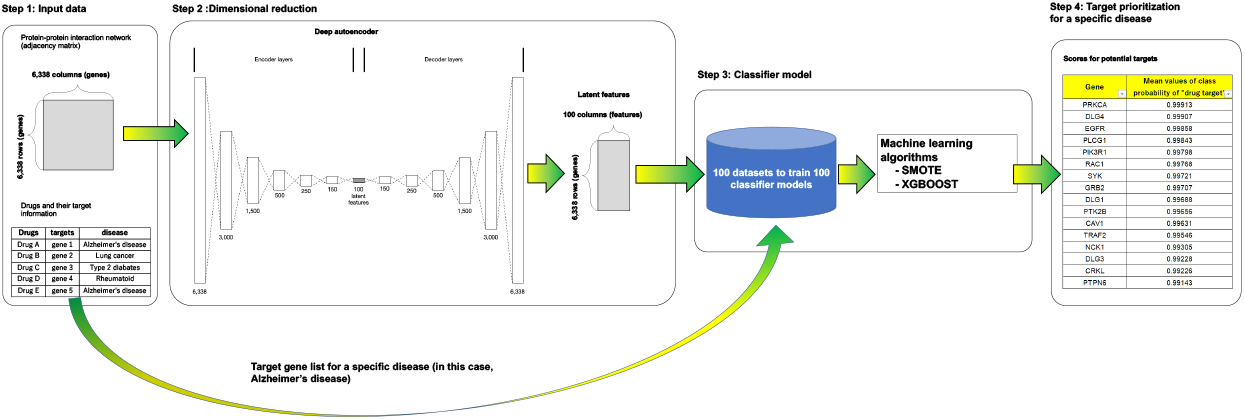
Computational analysis pipeline for drug target prioritization. **(Step 1)** Our computational framework used genome-wide protein-protein interaction networks and information of drug targets obtained from public domain databases. **(Step 2)** The framework is based on deep autoencoder to extract low-dimensional latent features from high-dimensional PIN. **(Step 3)** By using features from step 2 and target gene list for a specific disease, we build 100 datasets to train 100 classifier models. By using the 100 datasets and state of the art machine learning techniques (SMOTE and Xgboost), we build 100 classifier models to infer potential drug targets. **(Step 4)** We applied the classifier models to all the non-known drug-target genes in the PIN for prioritizing potential drug target genes. See “materials and methods” and “results and discussions” for details.

### Network embedding: Deep autoencoder based dimensional reduction of protein interaction network

We obtained directed human PIN from [20] and the PIN is composed of 6,338 genes and 34,814 interactions (see Materials and Methods for details). We generated an adjacency matrix for the human PIN. Elements in the matrix are represented as binary value, i.e., 1 or 0 in position (i, j) represents whether protein *j* is downstream interacting partner of protein i or not. The resultant matrix is composed of 6,338 rows and 6,338 columns. Each row in the matrix presents interacting pattern for each gene and used as features of the gene. Because the number of genes in the PIN is 6,338, the features for each gene are of 6,338 dimensions., i.e., a gene is characterized by 6,338 dimensional features based the PIN data.

As shown in Figure 1, in order to map high dimensionality of features (6,338 dimensions) for each gene into low dimensional features, we built and used a deep autoencoder. The deep autoencoder is composed of 7 encoder layers (6338-3000-1500-500-250-150-100) and symmetric decoder layers (100-150-250-500-1500-3000-6338) (see Figure 1). In the deep autoencoder, layers are fully connected and weights of links connecting layers are optimized by minimizing binary cross-entropy loss between values of nodes in input layer and those in output layer (for details, see materials and methods). After the optimization, for each gene, we used the optimized deep autoencoder to map high dimensionality of original features (6,338 dimensional features) into low dimensionality (100 dimensional features) through the middle layer (layer with 100 nodes) in the network. The resultant features for each gene are of 100-dimensional features.

The low-dimensional latent space contains enough information required to represent original high-dimensional human PIN. However, it is still unclear whether the low-dimensional features in the latent scape can explain topological and statistical properties obtained from the representative network metrics. In order to examine this issue, we calculated 9 representative network metrics for each gene in the PIN (e.g., in_degree, out_degree, betweenness, closeness, page rank, cluster coefficient, nearest neighbour degree (NND), bow-tie structure, and node dispensability, see methods for details) and compared the metrics with 100-dimensional features for the gene from the network embedding analysis (see Figure 2). As shown in the figure, among the 100-dimensional feature, several features are strongly correlated with out_degree, page rank, and closeness (*r* > 0.6, *r* indicates Spearman’s correlations between a feature and a network metric). Betweeness, in-degree, and bow-tie (input layer) are moderately correlated with several features (0.6 > *r* > 0.4), while NND and bow-tie (output layer and core layer) shows moderate negative correlations with several features (−0.6 > *r* > −0.4). In addition, cluster coefficient and node dispensability show weak correlation with several features (0.4 > *r* > 0.3). Interestingly, several features (e.g., dimensions 58, 86, 88, and 89) do not correlate with all of the 9 representative network-metrics. There results indicate that the low-dimensional features from network embedding analysis can capture the topological and statistical properties from network metrics. At the same time, the low-dimensional features from network embedding analysis may be able to capture information that are not obtained from analysis using representative network metrics.

**Figure 2.**
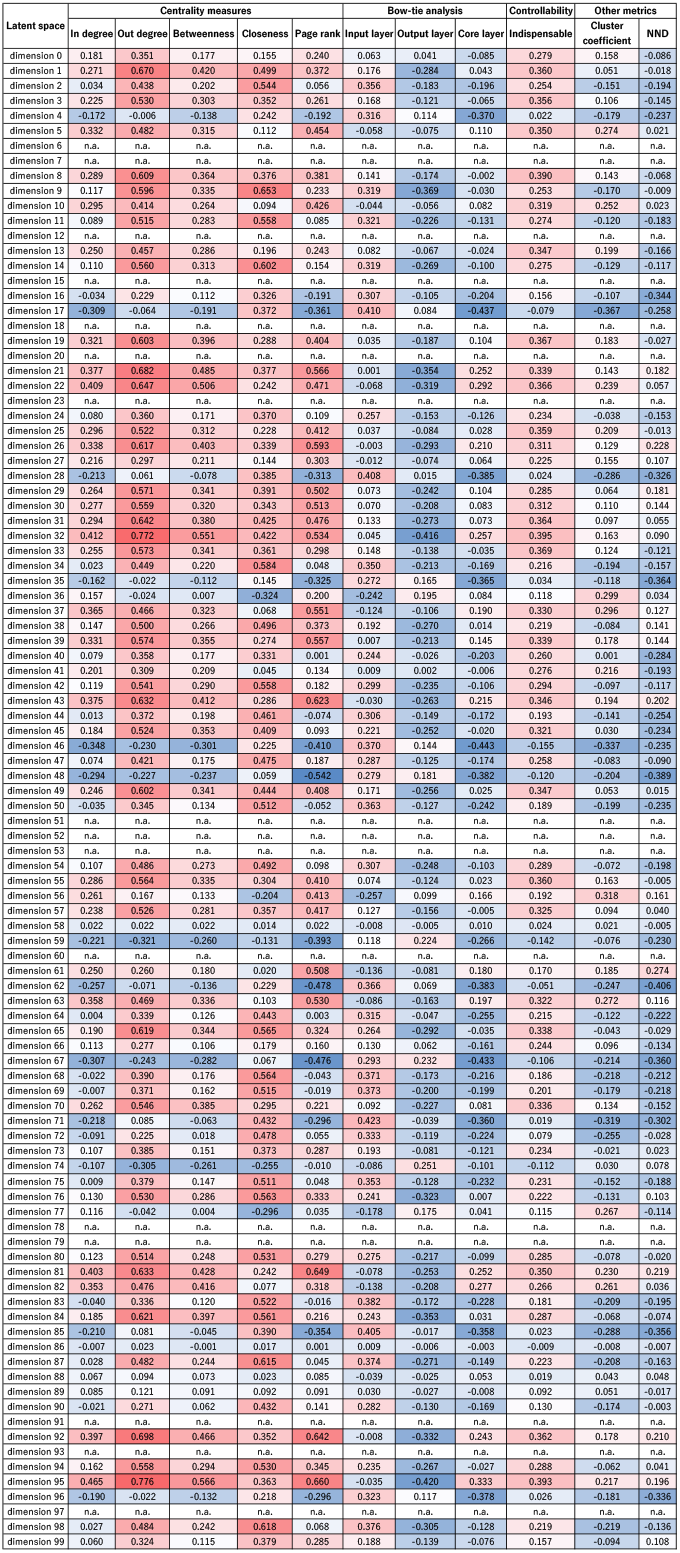
Relationships between features in low-dimensional latent space by deep autoencoder and representative network metrics in the PIN. Rows and colunms represent names of features in low-dimensional latent space and names of network metrics, respectively. The numeric in a cell represent Spearman’s correlation coefficient between a given low-dimensional feature and a given network metric, i.e., the correlation coefficient between feature “Dimension 1” and network metric “out_degree” is 0.67. Darker red (blue) indicate higher (lower) correlation coefficient.

### Machine learning based drug target prediction by using the extracted feature from the human protein network

In this study, we treated drug-target prediction problem as binary classification problem. In order to build binary classifier for drug-target prediction, we generated a training dataset by using the low-dimensional features extracted from the PIN and public domain drug-target information. From the public domain drug-target database, we obtained known drug-target genes for Alzheimer’s disease. Among the known targets, we could map 31 targets on the PIN. We regarded these 31 genes as positive cases, while we selected negative cases from remaining 6,307 (non-known target) genes. We randomly selected 500 negative cases (genes) among the 6,307 genes 100 times to build 100 datasets composed of 500 negative and 31 positive cases (genes). In the 100 datasets, each gene has 100 dimensional features that were obtained from deep autoencoder. We used the 100 datasets to build 100 binary classifier models to predict novel candidate targets for Alzheimer’s diseases.

The 100 datasets are class-imbalanced (e.g., 31 and 500 positive and negative cases, respectively) and classification using class-imbalanced data is biased in favour of the majority class. Further, in the datasets, the number of “positive#x201D; cases are too small, i.e., there are only 31 positive cases in the datasets. These problems can be attenuated by using over-samplings that are often used to produce class-balanced training datasets from class-imbalance data. In order to make class-balanced training datasets for building binary classifiers, we used a state of the art sampling method, SMOTE (Synthetic Minority Oversampling TEchnique) [21] that synthetically creates new cases in minority class (in this study, “positive” case) (see Materials and Method in details).

By using the class-balanced training datasets from SMOTE, we trained binary classifiers for drug target prediction. The binary classifier models are based on, Xgboost algorithm, the most efficient implementation of gradient boosting algorithm [22]. The trained binary classifier models calculate two class probabilities for each gene based on 100 dimensional features for each gene (e.g., probability of positive and that of negative), i.e., a gene with higher class probability of “positive” is more likely to be a member of “positive” class.

In order to optimize the binary classifiers based on Xgboost for drug target prediction, we conducted grid search with 5-fold cross validations. Please note that, in order to avoid data leakage, we conducted data splits for cross validations before SMOTE based over-sampling to generate class balancing training datasets. In order to evaluate predictive performance for each parameter combination, we calculated area under the receiver operator characteristic curve (AUC ROC). The mean value of AUC ROC for the 100 binary classifiers with optimal parameters is 0.648. It indicates that the 100 binary classifiers tend to assign high class probability of positive for known drug-target genes for Alzheimer’s disease. Therefore, non-known drug-target genes with high probability of “positive” may be potential novel drug-targets for Alzheimer’s disease.

We used the 100 trained binary classifiers to calculate class probability of positive and that of “negative “for all of the 6,307 non-known drug-target genes in the PIN. We used the mean value of class probability of positive from the 100 binary classifier to prioritize the 6,307 genes to infer putative therapeutic targets for Alzheimer’s disease (see Table 1 and Supplementary Table 1 for details), i.e., non-known targets with higher mean value of class probability of positive (e.g., DLG4 in Table 1 and Supplementary Table 1) may be more likely to be potential novel drug targets. 201 non-known drug-target genes showed mean value of class probability of “positive” higher than 0.75 (see Supplementary Table 1). We regarded these 202 genes as putative novel targets genes for Alzheimer’s disease.

**Table 1.**
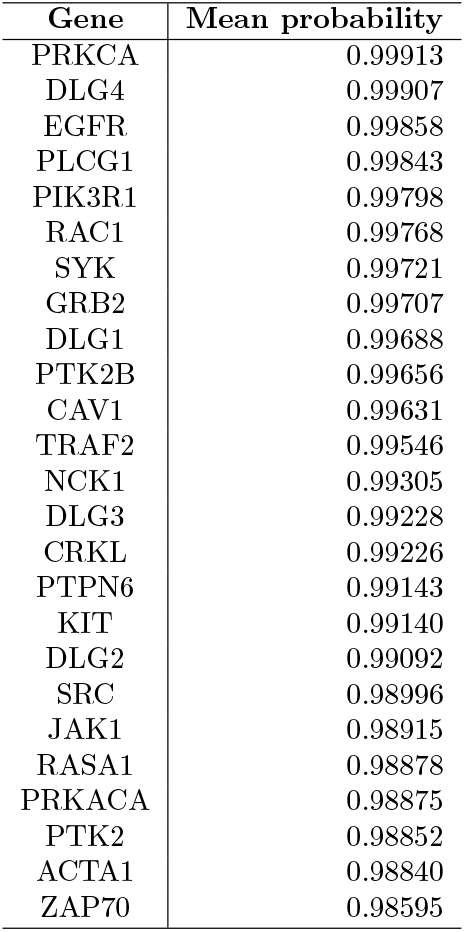
Top 20 genes with highest mean values of probability of “positive (drug target)” class.

### Pathway enrichment analysis

In order to infer potential target pathways for Alzheimer’s disease, we investigated significant pathways associated with putative 201 targets inferred by our computational framework (see Figures 3, 4, and 5). The 201 putative targets were significantly associated with pathways that control Alzheimer’s disease mechanisms (e.g., cytokine related signalling pathways, EGF receptor signaling pathway). Especially, among the significant pathways, those associated with inflammation mechanisms and immune systems. Especially, innate immune system is key components of Alzheimer’s disease pathology [23], i.e., continuous amyloid-*β* formation and deposition chronically activate immune system, causing disruption of microglial clearance systems [23]. These results indicated that we may be able to supress progression of Alzheimer’s disease by modulating these pathways, especially immune system and inflammation related pathways, through targeting these putative target genes.

**Figure 3.**
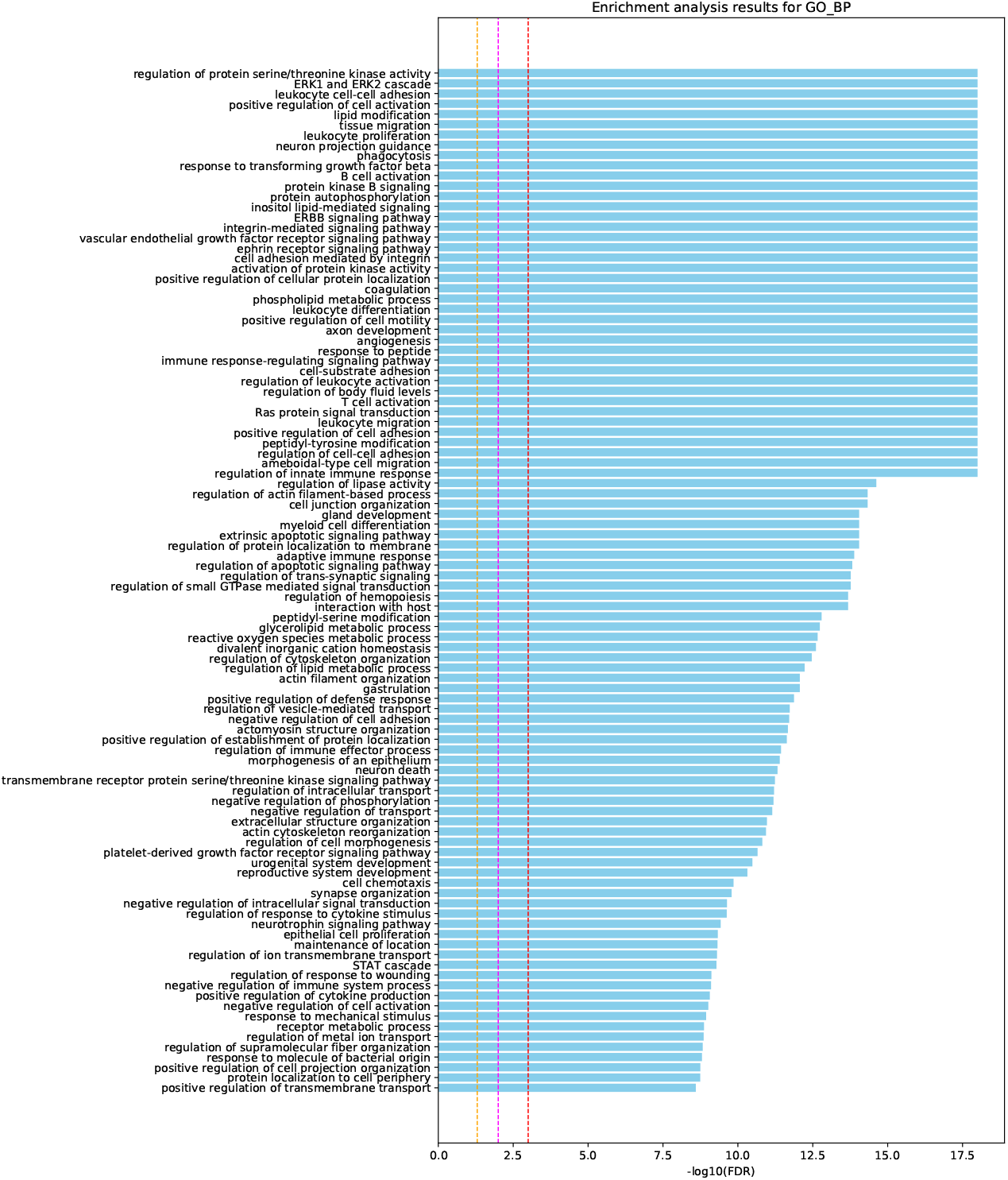
Pathway enrichment analysis with GO biological database for 201 putative targets for Alzheimer’s disease obtained from our computational pipeline. The names of pathways are shown on the vertical axis, and the bars on the horizontal axis represent the — log_10_(*p – value*) of the corresponding pathway. Dashed lines in orange, magenta, and red colors indicate p-value ¡ 0.05, 0.01, and 0.001, respectively.

**Figure 4.**
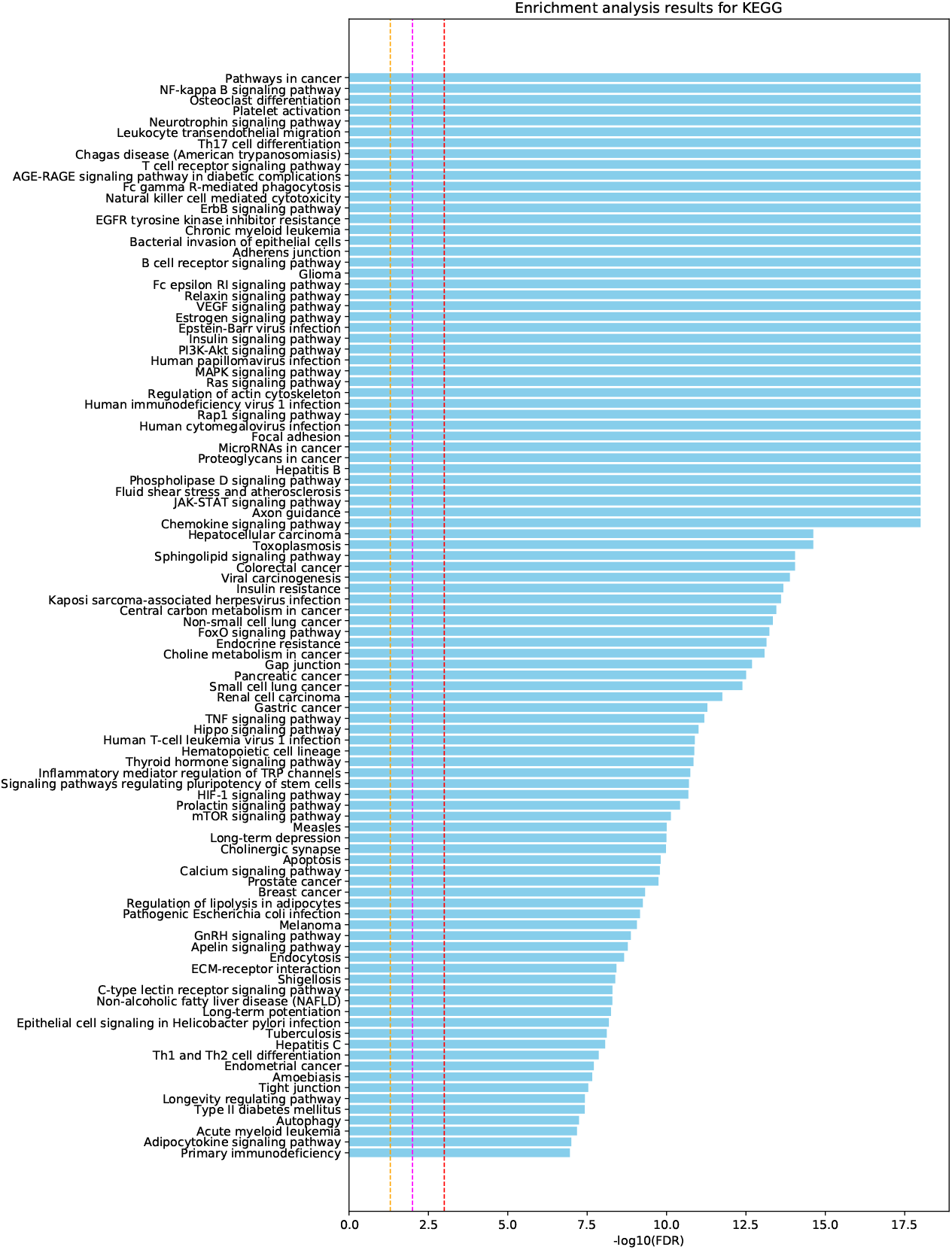
Pathway enrichment analysis with KEGG database for 201 putative targets. The legends for the figure are the same as that for Figure 3.

**Figure 5.**
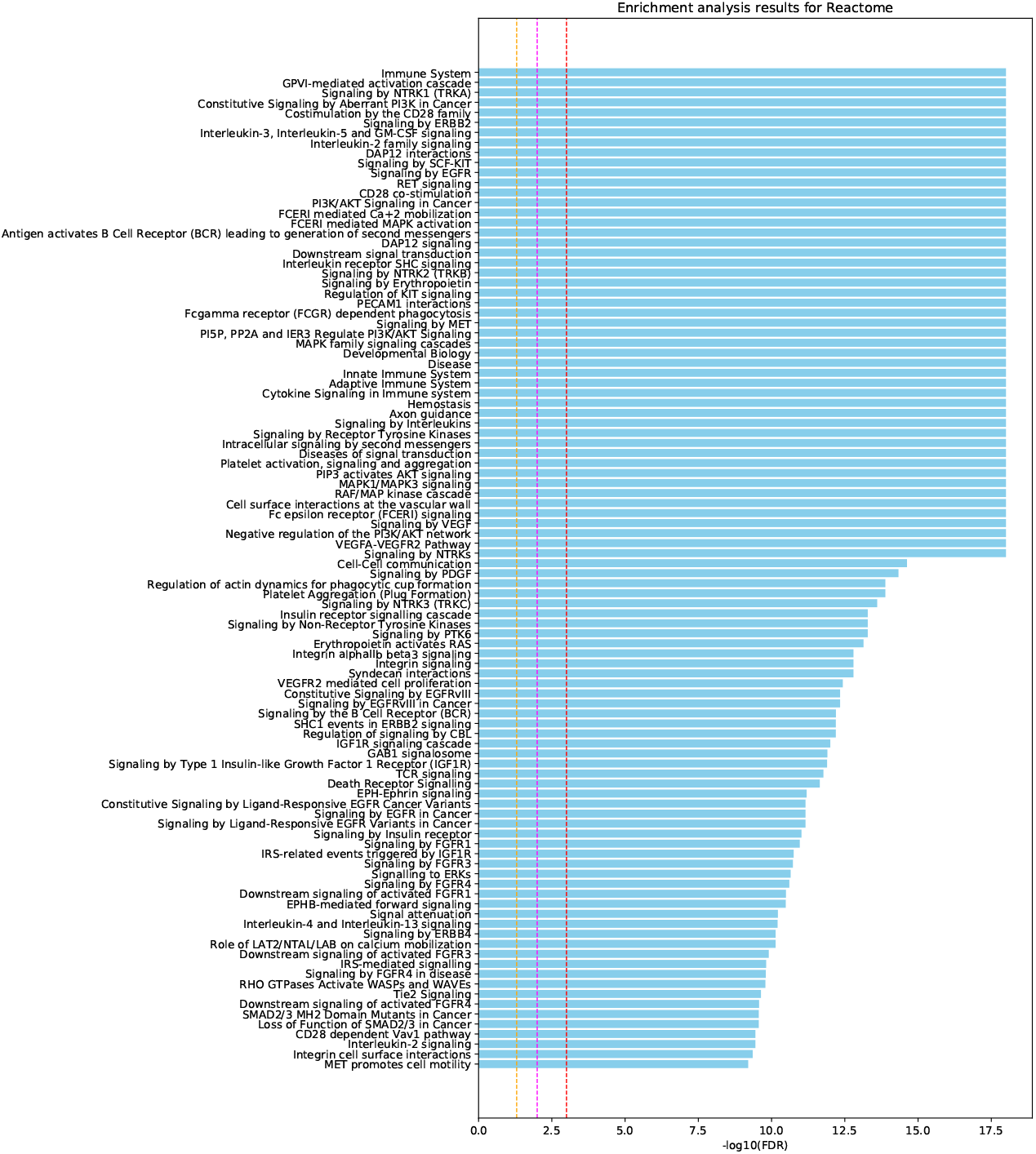
Pathway enrichment analysis with Reactome pathway for 201 putative targets. The legends for the figure are the same as that for Figure 3.

### Putative targets from our computational framework

Among 201 putative targets from our analysis (see Supplementary Table 1), we investigated top ranked genes and found that several top ranked genes play an important role in disease mechanism of Alzheimer’s disease.

For example, 2nd ranked putative target, DLG4 encodes, PSD95, a key protein for synaptic plasticity and down-regulated under aged and Alzheimer’s disease patients. Recently, Bustos et al demonstrated that epigenetic editing of DLG4/PSD95 ameliorate cognitions in model mice with Alzheimer’s disease [24]. Thus, epigenetic editing of DLG4 may provide a potential novel therapy to rescue cognitive impairment of Alzheimer’s disease.

The third ranked putative target is EGFR that is frequently upregulated in certain cancers. Wang et al. demonstrated that upregulation of EGFR cause memory impairment in amyloid-*β*-expressing fruit fly model [25]. Furthermore, they administrated several EGFR inhibitors (e.g., erlotinib and gefitinib) to transgenic fly and mouse model for Alzheimer’s disease and found that the inhibitors prevent memory loss in the two animal models. Based on the observations, they suggested that EGFR may be a potential therapeutic target to treat amyloid-*β* caused memory impairment.

The sixth ranked putative target is Rac1, a small signalling GTPase, that controls various cellular processes including cell growth, cellular plasticity, and inflammatory responses. Inhibition of RAC1 down-regulates amyloid precursor protein (APP) and amyloid-*β* through regulation of APP gene in hippocampal primary neurons [26]. RAC1 inhibitors can also prevent cell death caused by amyloid-*β*42 in primary neurons of hippocampus and those of entorhinal cortex [27]. Furthermore, based on analysis of protein-domain interaction network together with experiments using drosophila genetic models, Kikuchi et al. demonstrated that RAC1 is a hub gene in the network and thus causes age-related alterations in behaviour and neuronal degenerations [28]. Thus, RAC1 gene may be a potential therapeutic target to prevent amyloid-*β* induced neuronal cell death in Alzheimer’s disease.

The seventh ranked potential target is SYK, Spleen Tyrosine Kinase, that have potential to modulate accumulation of amyloid-*β* and hyperphosphorylation pf Tau associated with Alzheimer’s disease [29]. Nilvadipine, an antagonist of L-type calcium channel (LCC), inhibits accumulation of amyloid-*β*, but this is not due to LCC inhibition, but to other mechanisms. Paris et al. demonstrated that down-regulation of SYK exert similar effect of (-)-nilvadipine enantiomer on clearance of Abera and reduction of Tau hyperphosphorylation [29]. Schweig et al. demonstrated that, in mices with overexpression of amyloid-*β*, SYK activation occurred in microglia and increased neurite degeneration due to amyloid-*β* plaques associated with aging [30]. They also demonstrated that, in those with overexpression of Tau, SKY activated in microglia and misfolded and hyperphosphorylated tau accumulated in hippocampus and cortex. Furthermore, Schweig et al. demonstrated that, by immunoprecipitation and RT-PCR experiments, SYK inhibition induces reduction of Tau in an autophagic manner [31]. They also showed that SYK acts as an upstream target in the mTOR pathway and SYK inhibition induces Tau degradation through decreasing mTOR pathway activation.

The 10th ranked putative target, PTK2B, is a key gene to mediate synaptic dysfunction induced by amyloid-*β* in Alzheimer’s disease [32]. Salazar et al. demonstrated that, in transgenic mice model of Alzheimer’s disease, PTK2B deletion improves deficits in memory and learning functions as well as synaptic loss [32].

In addition, although SOCS1 is the 86th ranked putative targets, SOCS1 modulates cytokine responses through suppression of JAL/STAT signaling to control CNS (central nerve system) inflammation [33]. Thus, SOCS1 may be a potential key therapeutic modulator in disease state of Alzheimer’s disease.

These observations indicated that our computational framework successfully identified key genes that may be novel target candidates for Alzheimer’s disease.

### Inference of repositionable drug candidates

Drug repositioning is to apply an existing drug for a new indication that is different from original indication. The advantage of drug repositioning is the established safety, i.e., studies of toxicology have been already done on a target drug. Therefore, development of computational methods to predict repositionable candidates could be promising strategy to reduce the cost and time that are inevitable for drug development.

Researches have proposed various drug repositioning methods. We can roughly classify these methods into two different major categories, Activity-based drug repositioning and in silico drug repositioning. With the former approach, a number of drugs for non-cancerous diseases are discovered for cancer therapeutics [34]. In recent years, the latter approach becomes successful because of the enhancement of the protein-protein interaction database, protein structural database and in-silico network analysis technology. Such kind of applications about drug repositioning via network theory are discussed. Iorioet et al [35] reported that Fasudil (a Rho-kinase inhibitor) might be applicable to several neurodegenerative disorders, by verifying similarity between CDK2 inhibitors and Topoisomerase inhibitors. Cheng et al. [36] applied three similarities (drug-based, target-based, and network-based similarities) based inference methods to predict interactions between drugs and targets, and finally confirmed that five old drugs could be repositioned.

As discussed in the precious paragraph, in-silico network based approaches may be the most promising tools towards computational drug repositioning. Especially, networks connecting drugs, targets, and diseases could be useful resources to investigate novel indications for FDA-approved drugs, i.e., if a target gene P is a putative target for a disease A and is a known target gene of drug R for a different disease B, the disease A may be a potential novel target disease for the drug R (see Figure 6). Thus, in order to infer potential repositionable drugs and their potential target disease, we further investigate the list of 201 predicted putative target genes (gene with class probability of target class > 0.75 in Supplementary Table 1) from our computational framework and drug-target information across different diseases, i.e., if at least one targets of an existing drug are among 201 putative targets, we regard the drug as potential repositionable drug. As shown in Supplementary Table 2, we inferred 332 candidate repositionable drugs for Alzheimer’s disease. For each candidate repositionable drug, we calculated the number of overlapped genes between know targets of the drug and 201 putative targets. We ranked candidate repositionable drugs based on the number of overlapped genes. Among the predicted repositionable drug candidate, top ranked candidates may have efficacy for the target disease. Table 2 listed the top 20 highest ranked candidate compounds.

**Figure 6.**
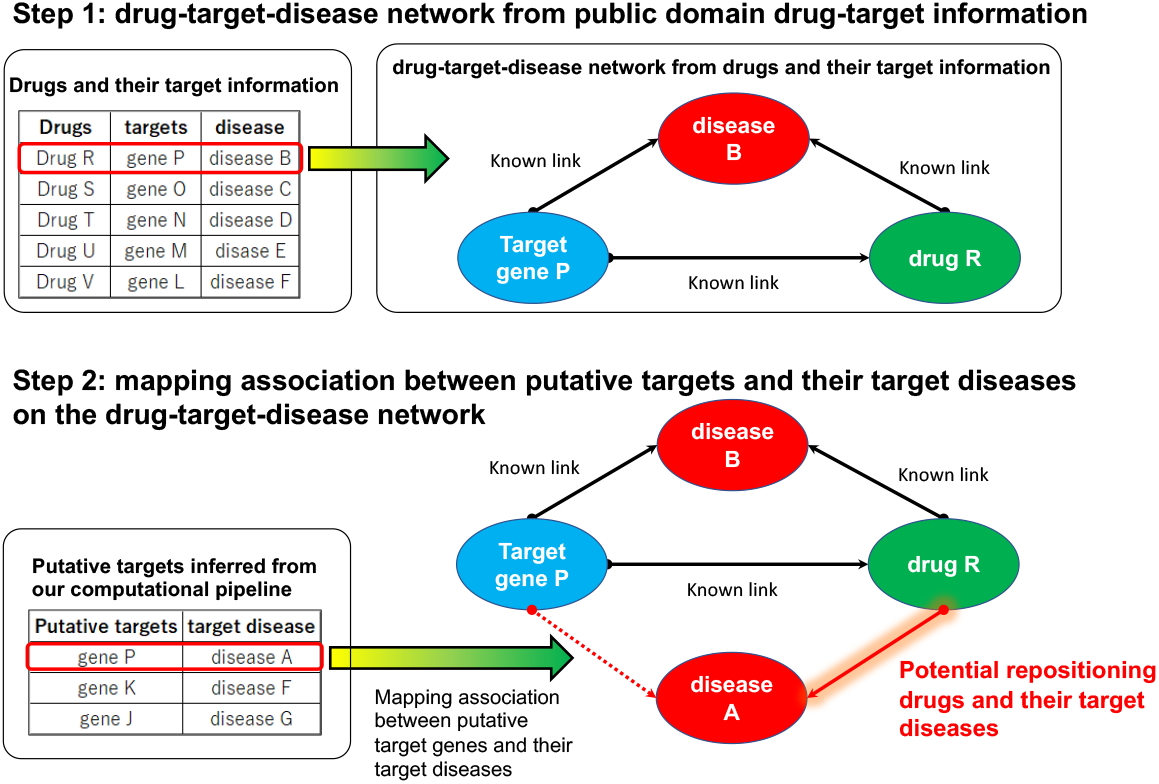
A method to infer potential repositionable drugs based on putative targets from our computational pipeline. Step 1: We obtained drug-target-disease network from Drug-Bank database. Step 2: We mapped associations between putative target genes and their target diseases to infer potential repositionable drugs for a given disease.

**Table 2.**
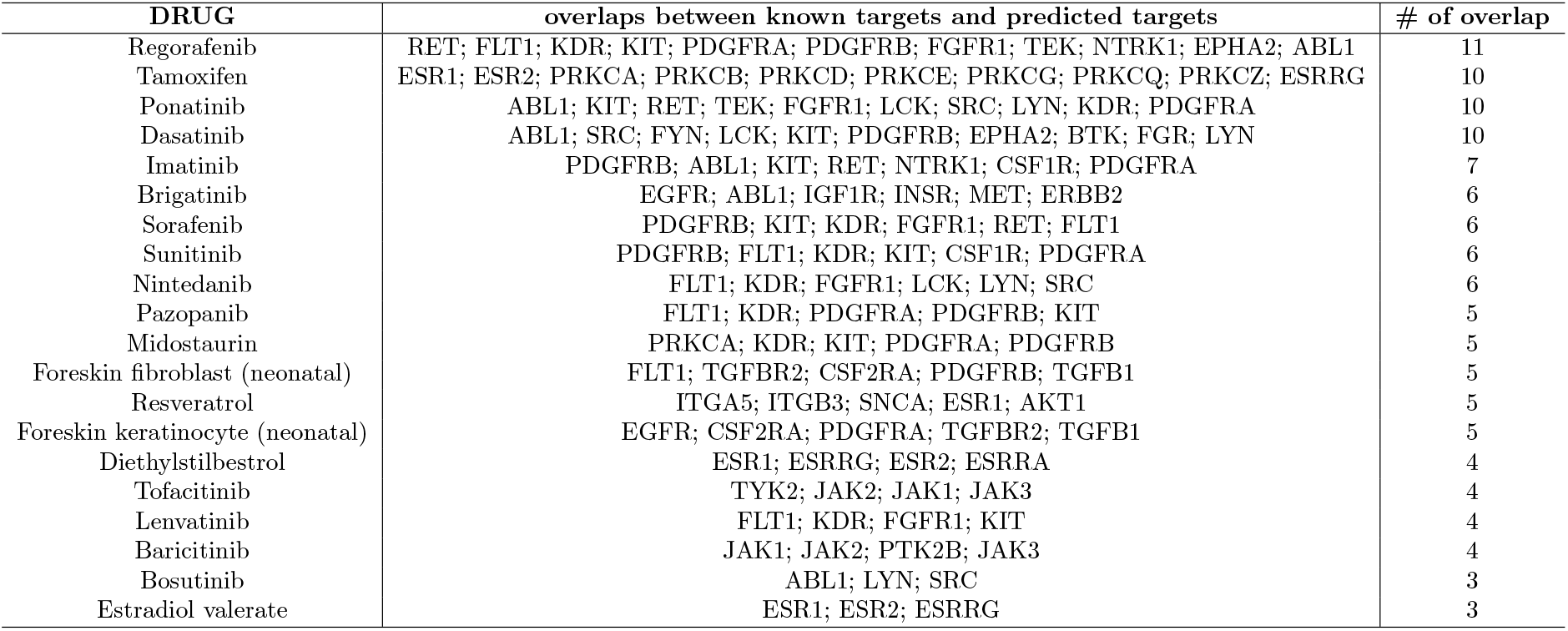
Top 20 ranked candidate repositioning drugs for Alzheimer’s disease

For example, our method predicted that Tamoxifen (top 2rd ranked candidate), a FDA-approved estrogen receptor modulator to treat hormone-receptor-positive breast cancer patients, as a potential drug target for Alzheimer’s disease. As mentioned in Wise PM [37], estrogens therapy could protect neuronal cells against cell death through modulating expression of genes that are keys to inhibit apoptotic cell death pathways. Indeed, based on nation-wide cohort study in Taiwan, Sun et al. reported that patients with long-term use of tamoxifen exhibit reduced risk of dementia [38].

Our method also predicted Bosutinib (top 20th ranked target), a FDA-approved tyrosine-kinase-inhibitor (TKI) drug (Bcr-Abl kinase inhibitor) to treat Philadelphia chromosome-positive (Ph+) chronic myelogenous leukemia, may be a potential repositionable drug for Alzheimer’s disease (see Table 2). Lonskaya et al reported that Bosutinib together with Nilotinib systematically modulate immune system in CNS through inhibition of non-receptor tyrosine kinase Abl to clear out amyloid and to decrease neuro-inflammation [39]. It indicated that TKIs, especially, Bosutinib could be potential repositionable drugs to treat early stage of Alzheimer’s disease.

Among the predicted repositionable candidates, 23 are immunosuppressive agents. The 23 candidates may include promising repositionable drugs for Alzheimer’s disease, because immune mediated inflammation in central nerve systems play an important role in disease mechanisms of Alzheimer’s disease. Among the 23 candidates, Dasatinib (4th ranked compound) may be the most promising candidate. Recently, Zhang et al reported that senolytic therapy (combinatorial drug therapy of dasanitib together with quercetin) has potential to reduce production of proinflammatory cytokine and to alleviate deficits of cognitive functions in Alzheimer’s disease mouse models, through selective removal of senescent oligodendrocyte progenitor cells [40,41]. Furthermore, Dasatinib plus quercetin is now registered in a clinical trial (ClinicalTrials.gov Identifier: NCT04063124).

These observations suggested that our method could be a powerful tool to infer potential repositionable drugs, especially for Alzheimer’s disease.

## Conclusion

In this study, we developed a deep autoencoder based computational framework that extracts low-dimensional latent space embedded in high-dimensional data of the human PIN and uses the features in the latent space to prioritize potential novel putative targets.

We examined relationships between the features in the latent space and the representative network metrics and found that the network metrics can explain a large number of features in the latent space, while the other features do not correlate with the network metrics. These results indicate that the features in latent space are likely to capture information that the representative network metrics can not capture, while the features also can capture information obtained from the network metrics.

We applied our computational framework to prioritized putative target genes for Alzheimer’s disease and successfully identified key genes (e.g., DLG4, EGFR, RAC1, SYK, PTK2B, SOCS1) associated with disease mechanisms of Alzheimer’s diseases. Furthermore, by using the putative targets from our computational framework, we successfully inferred promising repositionable candidate-compounds for Alzheimer’s disease (e.g., Tamoxifen, Bosutinib, Dasatinib).

It is pertinent to note here that our computational platform is easily applicable to investigate novel potential therapeutic targets and repositioning compounds for any diseases including rare diseases.

## Materials and Methods

### Protein-protein interaction network and drug-target information

We obtained directed protein interaction network from [20]. The network composed of 6,338 genes and 34,814 non-redundant interactions among the genes.

We obtained information of drugs and that of their target genes from DrugBank database [42] (http://www.drugbank.ca/). We manually investigated “description” field for all the drugs in the DrugBank database and identified 61 therapeutic drugs for Alzheimer’s disease. We regarded the 61 targets for the drugs as the established drug targets for Alzheimer’s disease. Among the 61 targets, 31 were mapped on the PIN.

### Feature extraction from PIN by Deep autoencoder

We build deep autoencoder with symmetric layer structure composed of 7 encoders layers and 7 decoder layers (e.g., 7 encoder layers (6338-3000-1500-500-250-150-100) and symmetric decoder layers (100-150-250-500-1500-3000-6338)). Layers are fully connected and layers except for output layer used rectified linear unit (ReLU) [43] as activation function. The output later used sigmoid function to make binary outputs. We optimized the deep autoencoder network by using “adam#x201D; [44] optimizer with learning rate of 1.0 × 10^-6^, the number of epochs = 10,000, batch size = 10, and default values for other parameters. In the optimization step, we minimize binary cross-entropy loss between values of nodes in input layer and those in output layer. We used a representative deep learning platform, Keras [45], with Thensorflow [46] backend to implement the deep autoencoder. To performe the deep autoencoder based dimensionality reduction analysis of PIN, we used Tesla K80 GPU on shirokane 5 super computer system (https://supcom.hgc.jp/english/).

### Statistical and topological analysis of the PIN

In order to investigate statistical topological features in the PIN, for each gene, we calculated representative network metrics, in_degree, out_degree, betweenness, closeness, page rank [47], cluster coefficient [48], nearest neighbour degree (NND) [49], bow-tie structures [50], and indispensable nodes [51, 52] in the PIN.

In degree; In_degree for a given node represents the number of nodes have link to the node (in other words, upstream neighbours of the node).

Out degree; Out_degree represents the number of links from the given node to other nodes (in other words, downstream neighbours of the nodes).

Betweenness; Betweenness for a given node i is the number of shortest paths between two other nodes that pass through the node i.

Closeness; The value of closeness for a given node i is the mean length of the shortest paths between the node i and all the other nodes in the network.

Page rank [47]; Page rank for a given node is a metric to roughly estimate the importance of the node in the network. The page rank score is calculated by the algorithm proposed by Google (see http://infolab.stanford.edu/~backrub/google.html for details of the algorithm). A given node has higher page rank, if nodes with higher rank have links to the node.

Cluster coefficient [48]; Cluster coefficient of a node *i* (*C_i_*) is calculated by using the following equation. 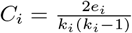, where *k_i_* is the degree of the node *i* and *e_i_* is the number of links connecting neighbour nodes of the node *i* to one another.

Nearest neighbour degree (NND) [49]; The value of NND for a given node *i* is the average degree among nearest neighbour nodes of the node *i*.

Bow-tie structure [50]; The biological networks often have bow-tie structures that are composed of three components (e.g., input, core, and output layers) [50]. Yang et al. proposed a bow-tie decomposition method to classify nodes in to three classes, nodes in input layer, those in core layer, and those in output layer [50]. In the decomposition analysis, a strongly connected component composed of the largest number of nodes is defined as the nodes in core layer. Nodes in input layers can reach the core layer, while those in core layer can not reach input layer. The nodes in core layer can reach the nodes in output layers, while nodes in output layers can not reach the core layer. We represent the analysis results from Bow-tie decomposition by one-hot vector encoding, i.e., we used three binary variables (variables for “input layer”, “core layer”, “output layer”) to represent the results from bow-tie structure. For example, for a node classified in to core layer, the value of core layer of the node is equal to 1, while the value of “input layer” and that of “output layer” is equal to be 0.

Indispensable nodes [51, 52]; Liu et al. developed a controllability analysis method to identify the minimum number of driver nodes (ND) that we must control to modulate dynamics of the entire network [52], i.e., they used the Hopcroft-Karp ‘maximum matching’ algorithm [53] to identify the minimum set of driver nodes [52]. Indispensable nodes that are potential key player nodes and are sensitive to structural changes in a network, are obtained from controllability analysis, i.e., removal of an indispensable node increase the ND in the network [51]. Vinayagam et al. reported that indispensable proteins in a human PIN tend to be targets of mutations associated with human diseases as well as those of human viruses [51]. We represent the analysis results of indispensable nodes by one-hot vector encoding, i.e., we used a binary variable to represent the results. For example, for an indispensable node, the value of binary variable of the node is equal to 1, while, for a non-indispensable node, the value is equal to be 0.

For the network analysis, we used igraph R package [54].

### Oversampling by SMOTE algorithm

In order to make class-balanced dataset for building binary classifier, We used a state of the art sampling method, SMORT [21] to generate class-balanced dataset to build binary classifier for drug target prediction. The SMOTE algorithm synthetically creates more cases in minority class. In order to synthetically generate cases in the minority class, the SMOTE algorithm selects k nearest neighbours of a case in minority class and randomly select a point along a line connecting them. The selected point is used as an additional case in the minority class. We used a python module,” imblearn”, to do oversampling based on SMOTE algorithm. We used *k* = 2 to do SMOTE based oversampling.

### Binary classifier model based on Xgboost

In order to build binary classifier for drug target prediction, we used Xgboost that is the most efficient implementation of gradient boosting algorithms [22]. The gradient tree boosting is among the state of the art supervised-learning algorithms. The algorithm makes a large number of weak learners and build a strong learner that is in the form of ensemble of the weak learners. In boosting step, the algorithm continues to update weak learners by correcting errors made by previous learners. After that, the algorithm aggregates the predictions from the weak learners to make the final prediction through minimizing the loss by using gradient descent algorithm.

To build Xgboost algorithm based binary classifiers, we used XGBClassifier and scikit-learn [55] python modules. The XGBClassifier has several parameters. We examined various values for each parameter (please see manual for XGBClassifier module, https://xgboost.readthedocs.io/en/latest/python/python_api.html, for details), learning_rate = (0.01, 0.1, 0.5), max_depth = (1, 2, 3, 5, 10), n_estimators = (100), gamma = (0, 0.3), boostor = (’gblinear’), objective = (’binary:logistic’), reg_lambda = (0, 0.1, 1.0), and reg_alpha = (0, 0.1, 1). For the other parameters, we used default value. To evaluate binary classifier models and optimize parameters of the models, we conducted 5-fold cross validation.

### pathway enrichment analysis

In order to identify significant pathways associated with putative targets inferred by our computational framework, we used WebGestalt web tool [56]. WebGestalt uses over-representation analysis (ORA) that statistically evaluates overlaps between gene set of interest and a pathway [57]. In the analysis, initially, the number of overlapped genes between the gene set of interest and a pathway is counted. Then, hyper-geometric test is used to examine whether the pathway is over- or under-representation in the gene set of interest (for each pathway, p-value and FDR is calculated based on overlap). Based on the ORA analysis, we examined the pathways in Reactome, Panther, KEGG, and GO biological processes and regarded the pathways with FDR ¡ 0.05 as significant pathways associated with the gene set of interest.

### source code availability

Documentation and source code are available at https://github.com/tsjshg/ai-drug-dev.

## Supplementary Information

`Supplementary_Table1.xlsx` and `Supplementary_Table2.xlsx` are available.

## Authors contribution

Conceived the experiments: ST, TH, AY, TN, SG, MK, SK, HK. Designed the experiments and analyses: ST, TH, AY. Performed the experiments: ST, TH. Analyzed the data: ST, TH, AY, TN. Wrote the paper: ST, TH, AY, TN, SG, MK. Supervised the research: ST, TH, HK, HA, HT.

## Acknowledgements

The authors would like to express their sincere gratitude to all individuals who have helped in this paper.

